# Sex-specific crossover rates did not change with parental age in *Arabidopsis*

**DOI:** 10.1101/2020.02.06.938183

**Authors:** Ramswaroop Saini, Amit Kumar Singh, Geoffrey J. Hyde, Ramamurthy Baskar

## Abstract

Crossing over, the exchange of DNA between the chromosomes during meiosis, contributes significantly to genetic variation. The rate of crossovers (CO) varies depending upon the taxon, population, age, external conditions, and also, sometimes, between the sexes, a phenomenon called heterochiasmy. In the model plant *Arabidopsis thaliana*, the male rate of crossovers (mCO) is typically nearly double the female rate (fCO). With increasing parental age, it has been reported that the disparity decreases, because fCO rises while mCO remains stable. That finding, however, is based on chromosome-based averaging, and it is unclear whether all parts of the genome respond similarly. We addressed this point by examining how the level of heterochiasmy responded to parental age in eight genomic intervals distributed across the five chromosomes of *Arabidopsis*. Unlike the previous work, in each of the eight intervals, the level of heterochiasmy did not change with age, that is, the ratio mCO:fCO remained stable. As expected, though, amongst the intervals, the levels of heterochiasmy at any of the four ages examined, did vary. We propose that while the levels of heterochiasmy in *Arabidopis* might decrease with age on a chromosomal basis, as reported earlier, this is not true for all locations within each chromosome. This has practical implications for plant breeding research, a major aim of which is identifying ways to induce local increases in CO rates.

## Introduction

During meiotic crossing over, homologous chromosomes align and exchange paternally and maternally derived DNA. Crossovers (CO) are one of the main sources of variation in sexually reproducing organisms, and as such, the rate at which they occur has considerable evolutionary significance (Ritz *et al.* 2017; Stapley *et al.* 2017). If the rate is too low, the organism has less chance of adaptation, if too high, an already effective genotype runs the risk of disruption. While the rate of crossovers can vary across taxa, populations, and between and within individuals, the possible scale of variation across these various levels appears remarkably constrained (Ritz *et al.* 2017). Nevertheless, the scope for some degree of CO rate variation exists for individual organisms, and is of practical importance, both medically and economically. For example, the frequencies of several forms of human chromosomal number abnormalities (in particular, trisomies) correlate with the increased frequency of CO (Hussin *et al.* 2011; Alves *et al.* 2017). In plant breeding, the development of ‘elite’ genotypes depends on meiotic COs that allow the accumulation of desirable traits, and much research is focused on finding ways to increase local CO rates (Wijnker and Dejong 2008; Crismani et al. 2013; Fernandes et al. 2018).

Interestingly, it is often not just the overall rate of CO that is important, but also the ratio of the male and female rates of CO (henceforth, mCO and fCO). In many taxa, these two rates differ to a greater or lesser extent, a phenomenon called heterochiasmy (Ritz *et al.* 2017; Stapley *et al.* 2017). At the most extreme, in what is called achiasmy, one sex does not form chiasmata at all (John *et al.* 2016; Satomura *et al.* 2019). This is the case,_for example, in the male of *Drosophila*, in which chromosome alignment employs an alternative to the synaptonemal complex (McKee *et al.* 2012). In true heterochiasmy, the ratio between the rates of the more and less recombinative sexes can vary from 1.035 to 14 (Ritz *et al.* 2017). Evidence indicates that, for a truly heterochiasmatic species, the sex that has the lower rate of CO will be the one for which genetic stability in the haploid phase is most likely to be critical to the future organism’s fitness (Lenormand 2003; Lenormand and Dutheil 2005; Stapley *et al.* 2017). In *Arabidopsis*, for example, its high self-pollination rate (95%; (Charlesworth and Vekemans 2005) suggests that the female haploid phase is most critical, thus possibly explaining why fCO has the lesser value (Lenormand 2003; Lenormand and Dutheil 2005). The ratio of mCO:fCO in young *Arabidopsis* seedlings is typically about 1.8 (Toyota *et al.* 2011; Giraut *et al.* 2011), but amongst different accessions, the values can vary by about 22% (López *et al.* 2012).

As well as having evolutionary drivers, both the overall, and sex-specific, CO rates, and also mCO:fCO, are influenced by age and extrinsic stressors such as temperature, pathogens, chemical exposure, and lack of nutrients (Hayman and Parsons 1962; Francis *et al.* 2007; Toyota *et al.* 2011; Hussin *et al.* 2011; Martin *et al.* 2015; Halldorsson *et al.* 2016; Li *et al.* 2017; Modliszewski and Copenhaver 2017; Saini *et al*. 2017; Stapley *et al.* 2017). The effect of age on mCO:fCO, and its mechanistic basis, has been much studied in humans because the increased rates of CO implicated in the chromosomal number abnormalities mentioned above mostly occur in older women (Hussin *et al.* 2011; Chiang *et al.* 2012; Nagaoka *et al.* 2012; Alves *et al.* 2017).

For plant CO, much less is known about age x sex interactions. For example, with respect to the influence of age on patterns of heterochiasmy in *Arabidopsis*, there has only been one study (Toyota *et al.* 2011); other studies have examined the response of mCO only (Francis *et al.* 2007; Li *et al.* 2017). In Toyota *et al.* (2011), the extent of heterochiasmy in primary shoots decreased with age, because, although there was no change in mCO, there was an increase in fCO. It is unclear, however, whether this pattern is true for each location in the chromosome. That study looked at 343 markers across the five chromosomes of the species, and reported on the average change of mCO and fCO for each chromosome, taking the mean of rates for the set of each chromosome’s applicable markers. The likelihood of intrachromosomal variation of heterochiasmic values is suggested by the results of Li *et al.* (2017). That study found that while mCO in primary shoots did not significantly change with age for markers in five of nine genomic intervals (thus in agreement with the earlier results of Toyota *et al.* 2011), the rates did significantly increase in two intervals. The possibility of intrachromosomal variation in *Arabidopsi*s heterochiasmy is also supported by other studies that have shown that, at least at one time point, the chromosomal average, and location-specific, values of mCO:fCO vary greatly depending on which chromosome is examined, and the location with the chromosome {Drouaud et al., 2007; Giraut et al., 2011}.

In this study we explore the possibility of intrachromosomal variation further, by looking at the influence of parental age on mCO, fCO, and mCO:fCO, using eight markers that cover all five chromosomes of *Arabidopsis*. Plants were sampled at four time points that cover the full reproductive duration of the *Arabidopsis* main shoot. We find that, while, at any one age, the ratio mCO:fCO differed both inter- and intra-chromosomally, the ratio, and also mCO and fCO, did not change with parental age of the main shoot. We believe the most likely reasons for the apparent discrepancy between our results and previous findings (i.e. Toyota et al., 2011) is that: (1) on the one hand, our small set of markers did not include any of the locations that exhibit an age-response by fCO nor, for that matter, any of those that exhibit an age-response by mCO (as reported in Li, 2017); (2) on the other hand, by reporting only on the chromosomal averages of mCO:fCO, the earlier study could not detect any of an intrachromosomal spectrum of age-responses. It appears this spectrum is wide enough to include a lack of response at some locations, as we have reported for all three parameters, and as Li et al. (2017) report for mCO.

## Materials and methods

### Plant growth conditions

Freshly harvested *Arabidopsis* seeds from Columbia or detector lines (described below) were surface sterilized with 70% ethanol, followed by 0.5% bleach treatment for 3 min. Subsequently, the seeds were washed thrice with sterile water and plated on autoclaved Murashige and Skoog media (MS, with 3% sucrose), pH 5.7, containing 0.05% Plant Preservative Mixture (Biogenuix Medsystem Pvt. Ltd., New Delhi, India) and incubated at 4° C in dark conditions, for synchronized germination. After 48 h, the plates were shifted to a seed germination chamber, with a uniform light intensity of 8000 lux units (16-h light/8-h dark cycle). The temperature of the chamber (Percival CU-36L6) was maintained at 22° C with a constant humidity of 80%. Three-week old seedlings were transferred from MS plates to soil and grown inside a plant growth chamber (Percival AR-36L3). The soil had equal proportions of garden soil, peat, perlite, and vermiculite (Keltech Energies Ltd., Bangalore, India).

### Arabidopsis detector lines used to score CO rates

To score CO rates, eight different detector lines covering at least one marker in each of the five chromosomes were used. The detector lines Col3-4/20, 3158 and 3162 were kind gifts from Avraham A. Levy (Department of Plant Sciences, Weizmann Institute of Science, Israel), (Melamed-Bessudo *et al.* 2005). Another set of detectors, the traffic lines CTL1.2, CTL1.18, CTL2.4, CTL4.7 and CTL5.17 were obtained from the Arabidopsis Biological Resource Center (Ohio State University, USA), (Wu *et al.* 2015) (Table 1). In all the lines, the eGFP and dsRed markers are driven by a seed-specific napin promoter. The detector lines, homozygous for both markers were crossed with Columbia plants, and the seeds obtained (heterozygous for both eGFP and dsRed) were used in the subsequent experiments.

**Table 1.**
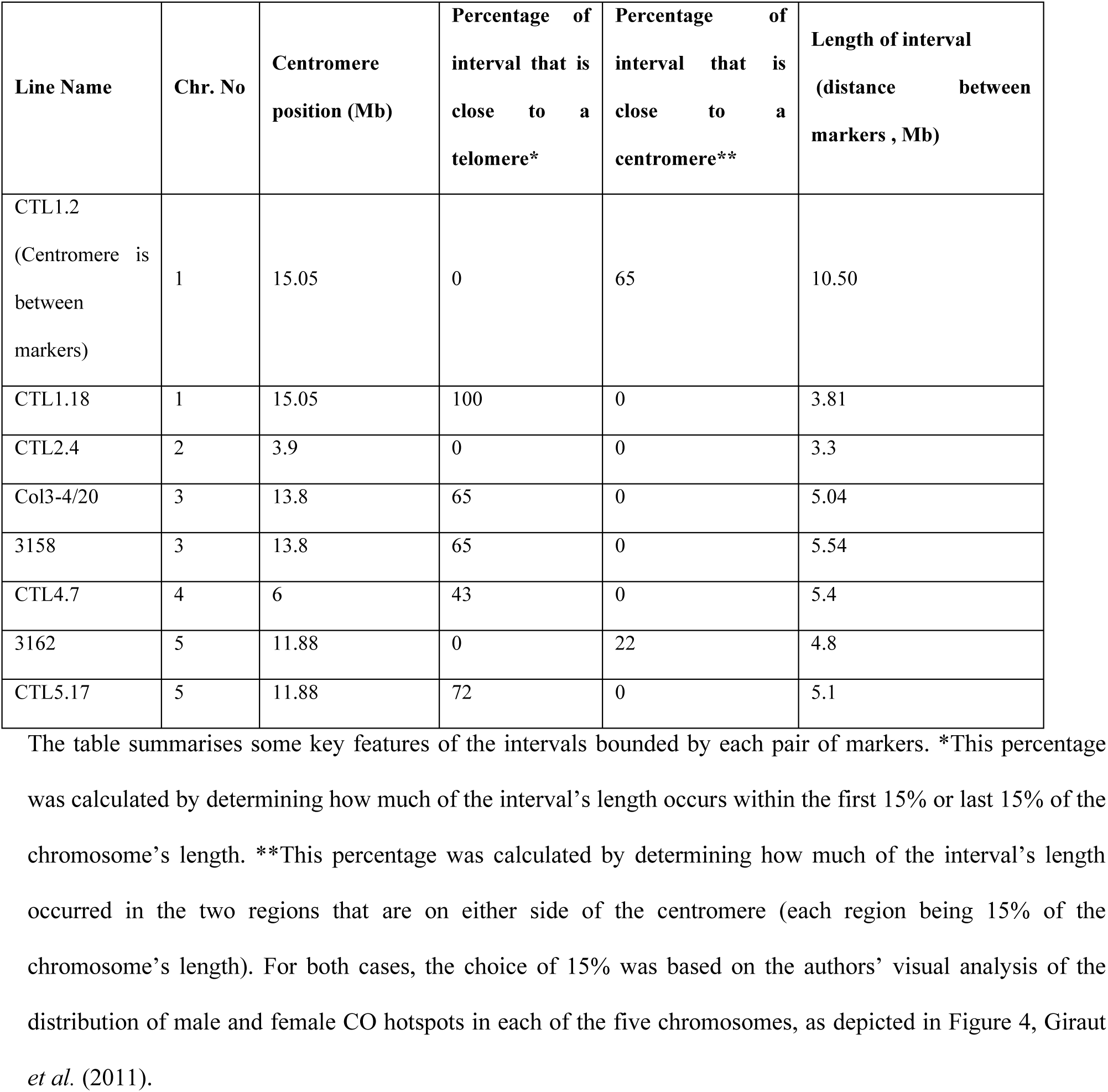
Structural features of the eight intervals used in this study.

### Investigating parental age effect on CO rates

To examine the influence of parental age on CO rates, plants of the detector lines and Columbia plants, of four different ages (40, 45, 50 and 55 DAS (days after sowing), were emasculated 48 h before pollination and reciprocally crossed with each other. Different colored threads were used to mark emasculated and pollinated flowers of different age groups. For each age, approximately 20 to 30 crosses were performed in three independent replicates. To score recombination during megaspore formation (fCO), we used emasculated flowers from the detector lines and crossed them with pollen from Columbia plants. Similarly, to estimate recombination rates during microspore formation (mCO), we used a detector line as the pollen donor for emasculated flowers of Columbia.

### Calculation of CO rates

The segregation of eGFP and dsRed markers (an indication of CO rates during micro- or mega-sporogenesis in the detector line parent), was analysed by the manual counting of seeds. Seeds were placed on a glass slide and analyzed under a Nikon Stereozoom Microscope (SMZ 1000) equipped with filters specific for both eGFP and dsRed. Images were captured for eGFP and dsRed separately and then both the images merged to identify the recombinant and non-recombinant seeds. CO rates were estimated based on the segregation of eGFP and dsRed markers. Of the four types of seeds obtained, seeds that fluoresce only either red or only green were counted as having undergone a CO, while the seeds that fluoresce for both red and green as well as those that do not fluoresce at all, were counted as not having undergone a CO. MR rates were calculated based on the formula:

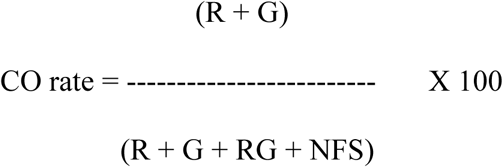

R- dsRed-only expressing seeds; G- eGFP only expressing seeds; RG- Seeds expressing both dsRed and eGFP; NFS- non-fluorescent seeds. The ‘rate’ is actually more correctly called a frequency, but we have used the term ‘rate’ because of its common usage in the literature.

### Statistical analysis

Meiotic CO rates follow a normal distribution and hence, a Gaussian generalized linear model (GLM) with identity link function was used (Nelder and Wedderburn 1972). The linear predictors were either the different ages, or the sex, of the detector-line parent. In all GLMs, the data from groups were compared. Correction for multiple testing was done to maintain the family-wise error rate at 5% (Gabriel 1969). Therefore, the *P* values were adjusted with a single-step method that considered the joint multivariate *t* distribution of the individual test statistic (Bretz *et al.* 2016). The results were reported with the two-sided *P* values adjusted for multiple comparisons (Singh *et al.* 2015). All statistical analyses were carried out in R (Team 2014). To adjust the *P* values for multiple testing, the R package multcomp was used with the test specification ‘single-step’(Bretz *et al.* 2016). Graphs were produced using GraphPad Prism 8.

## Results

### Heterochiasmy in eight intervals of Arabidopsis was unaffected by parental age

Using a set of eight *Arabidopsis* detector lines, we examined the influence of parental age on male and female CO rates. The eight detector lines heterozygous for both eGFP/dsRed were reciprocally crossed with Columbia wild type plants, with both parents being one of four ages (40, 45, 50, and 55 DAS) and MR rates were examined in the collected seeds (Figure 1). The eight intervals were distributed across all five chromosomes, and varied in length and the degree of overlap with sub-telomeric or pericentomeric regions (Figure 2; Table 1). One interval (in line CTL1.2) spanned the centromere.

**Figure 1.**
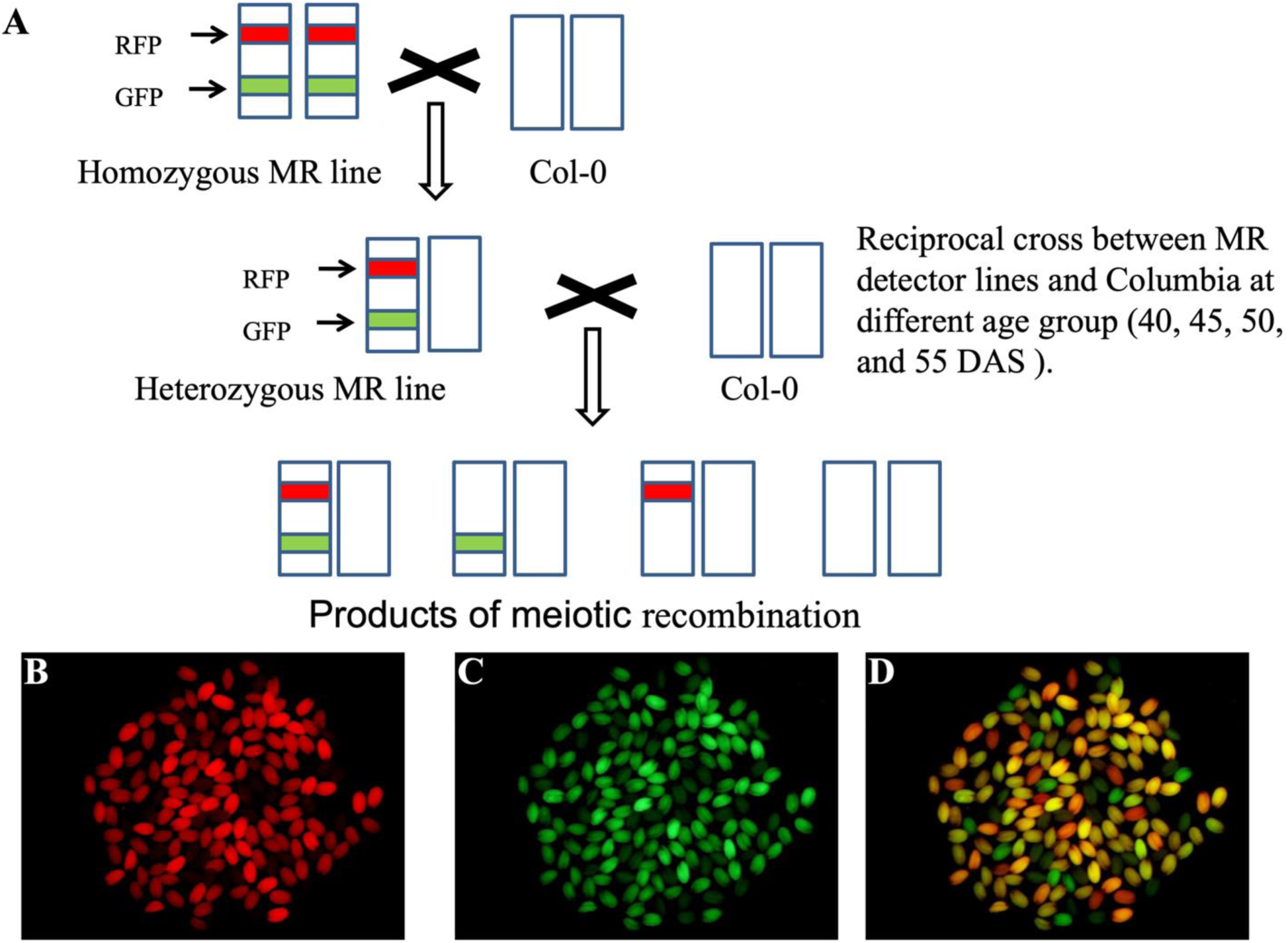
**(A) A** reciprocal cross between a heterozygous detector line and a Columbia plant results in seeds with one of four fluorescence patterns: a blend of red and green; only green; only red; no fluorescence. (**B**) A sample of seeds observed using a dsRed filter; (**C**) The same sample of seeds observed using an eGFP filter. (**D**) Merged image of B and C showing the four different patterns of fluorescence. Seeds in which a CO has occurred are those that either have only green or only red fluorescence, as determined by manual assessment of one image type or (if necessary) all three image types.

**Figure 1.**
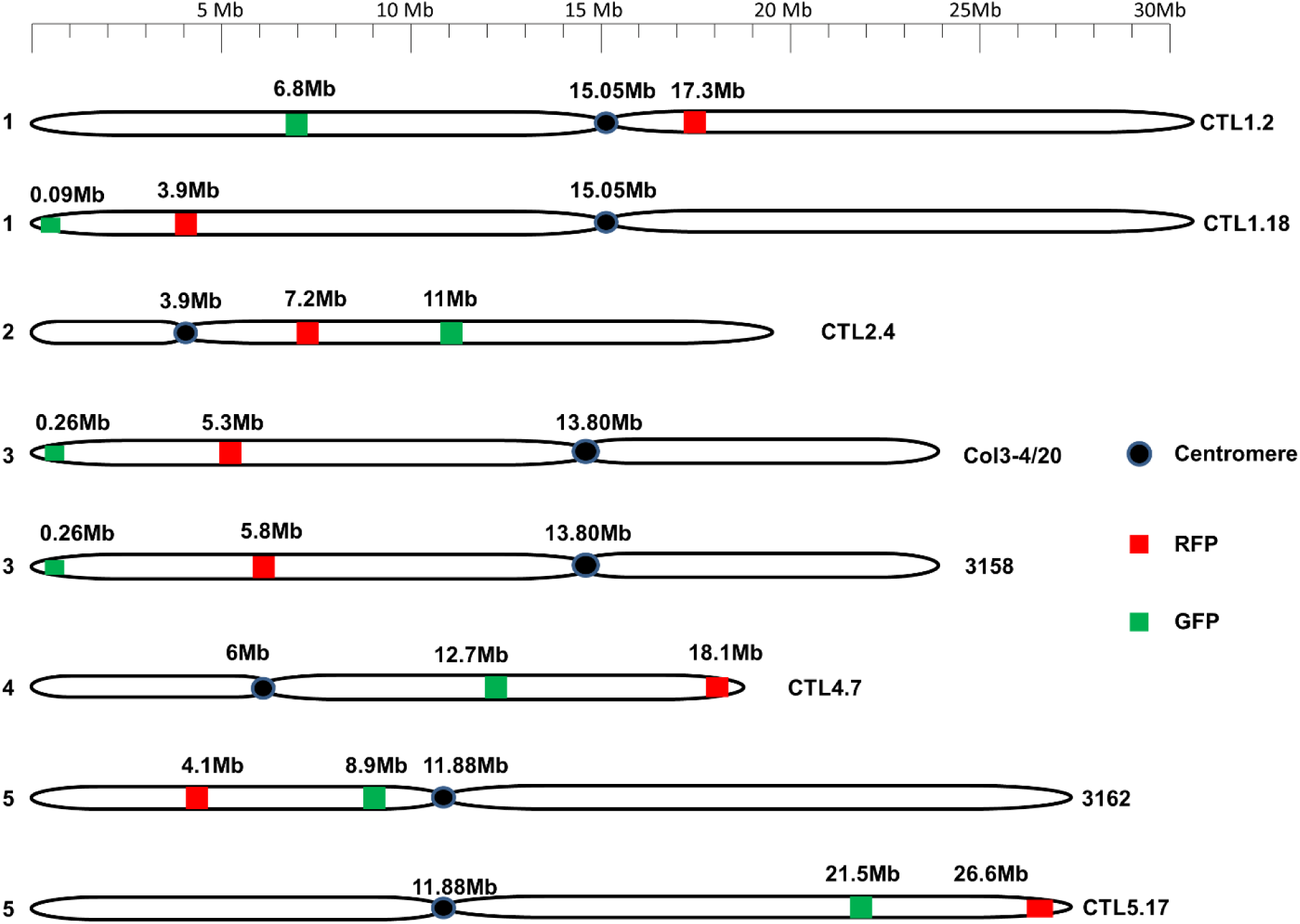
Physical maps of the chromosomes showing the location of the inter-marker intervals in the detector lines tested. Positions of eGFP and dsRed were drawn on the physical map using the chromosome map tool of The Arabidopsis Information Resource (TAIR).

For each of the eight intervals, there was no significant change in the ratio, mCO : fCO, as the age of the male and female parents was increased (Figure 3). Neither did the two individual rates that are used to calculate the ratio (i.e. mCO; and fCO) vary with age (Figure 3; Tables S1-8).

**Figure 3.**
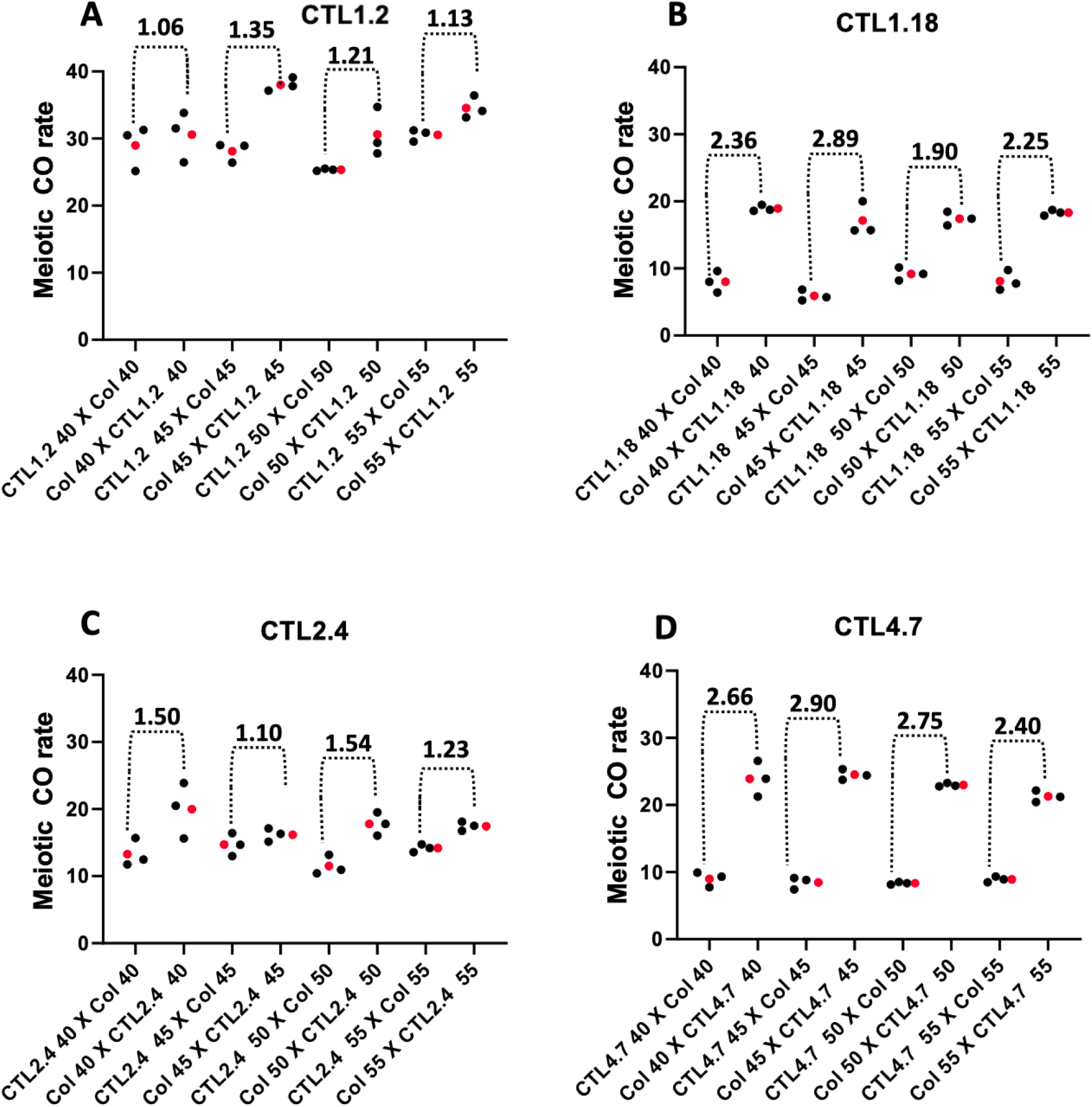
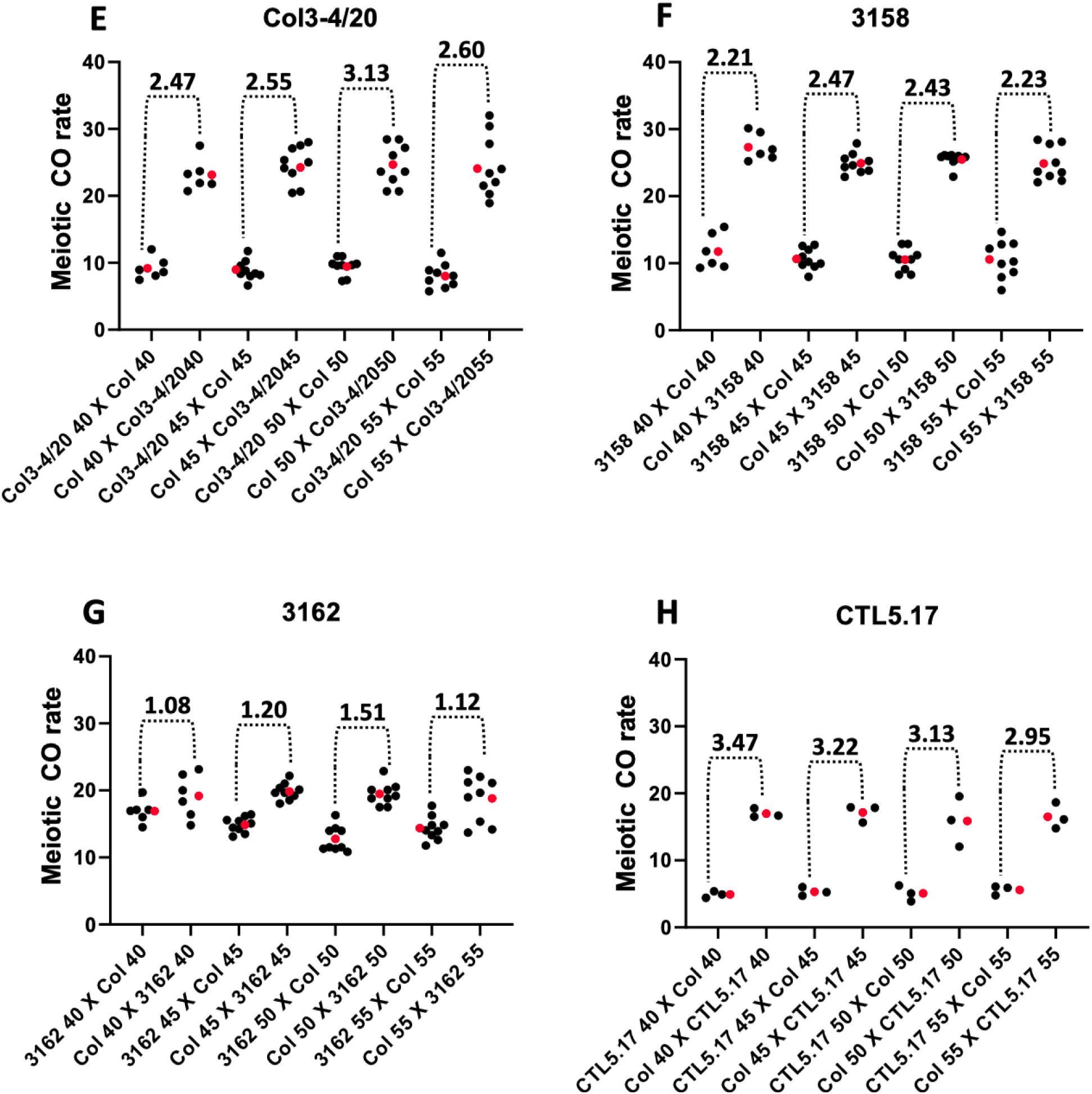
(A-H) Parental age did not affect CO rates or sex ratios: Reciprocal crosses between detector lines CTL1.2, CTL1.18, CTL2.4, CTL4.7, Col3-4/20, 3158, 3162, CTL5.17 and Columbia plants (of 40, 45, 50 and 55 DAS). Each dotted bracket spans two clusters of datapoints (comprising the full set of datapoints for one age category); the first cluster is of male CO rates, the second of female CO rates. The number above each bracket shows the average ratio of the male and female rates (i.e. mCO:fCO) for each age category. The graph represents individual replicated (black dots) and mean value (red dots) of the CO rates. GLM was used for detecting significant difference and *P* values were corrected for multiple testing (Supplementary Tables S1-8).

The average rates and thus the average heterochiasmic ratios, mCO:fCO, that we have measured can also be compared with those predicted by analysis of data published previously (Table S2 in Giraut *et al*. (2011)). Their genome-wide study reported the rates of mCO and fCO at 380 shared locations across all five *Arabidopsis* chromosomes, demonstrating remarkable variation in both rates from location to location. In Table 2 we present the measured and predicted rates and ratios (all ages combined), the predicted values being based on our interval-based analysis (Table S12) of the location-based data presented in their Table S2 (Giraut *et al.* 2011). The predicted values show a good congruence, in terms of their magnitudes relative to each other, with those that we measured. This point is evident when the measured and predicted ratios are ranked from high or low, according to their magnitude (Table 2).

**Table 2.** Measured and predicted male and female CO rates and their ratios.

| Line Name | Chr.<br>No. | Measured<br>average CO<br>rates in<br>males<br>(mCO) | Predicted<br>average CO<br>rates in<br>males<br>(mCO) | Measured<br>average CO<br>rates in<br>females<br>(fCO) | Predicted<br>average MR<br>rates in<br>females<br>(fCO) | Measured<br>mCO:fCO<br>(All ages) | Predicted<br>mCO:fCO |
| --- | --- | --- | --- | --- | --- | --- | --- |

|  |  |  |  |  |  | combined) |  |  |  |
| --- | --- | --- | --- | --- | --- | --- | --- | --- | --- |
|  |  |  |  |  |  | Ratio | Rank | Ratio | Rank |
| CTL1.2 | 1 | 33.44 | 45.26 | 28.24 | 37.59 | 1.184 | 7 | 1.20 | 8 |
| CTL1.18 | 1 | 17.95 | 19.43 | 7.81 | 5.98 | 2.30 | 4 | 3.25 | 2 |
| CTL2.4 | 2 | 17.84 | 17.84 | 13.41 | 13.41 | 1.33 | 5 | 2.31 | 6 |
| Col3-4/20 | 3 | 24.04 | 24.04 | 8.90 | 8.90 | 2.67 | 2 | 3.49 | 1 |
| 3158 | 3 | 25.66 | 25.66 | 10.88 | 10.88 | 2.33 | 3 | 3.06 | 4 |
| CTL4.7 | 4 | 23.17 | 23.17 | 8.67 | 8.67 | 2.67 | 2 | 3.21 | 3 |
| 3162 | 5 | 19.33 | 19.33 | 14.73 | 14.73 | 1.22 | 6 | 1.28 | 7 |
| CTL5.17 | 5 | 16.62 | 15.48 | 5.21 | 5.98 | 3.20 | 1 | 2.45 | 5 |

### Levels of heterochiasmy in eight intervals of Arabidopsis varied with the interval studied

The ratio mCO:fCO did, however, vary significantly on an interval by interval basis (Figure 3; Table 2). For example, the ratio of the average mCO:average mCO (across all four ages for any one interval) varied between 1.18 (CTL1.2) and 3.20 (CTL5.17; Table 2). These two lines also exhibited the most extreme values for both of the individual rates, mCO and fCO. CTL1.2, which had the lowest ratio, had the highest individual rates (mCO: 33.44; fCO: 28.24); while CTL5.17, which had the highest ratio, had the lowest rates (mCO: 16.62; fCO: 5.21) (Table 2).

The relatively higher values of the measured male and female CO rates of detector line CTL1.2 were also predicted by our analysis of the data from Table S2 of Giraut *et al.* (2011). A high CO rate in an interval can be the consequence of the number and strength of its CO hotspots and/or the length of the interval. In the case of CTL1.2, the high male rate was solely due to the interval being two to three times longer than any of the others. According to our analysis of Table S2, Giraut *et al.* (2011), the average male rate along this interval (as measured in the earlier study) is in fact the second lowest of the eight intervals. For the relatively higher female rate of CTL1.12, the interval’s female hotspots also contributed: it has the highest average rate of recombination of any of the intervals (calculated from Table S2, Giraut *et al.* (2011)).

The pattern of high or low sex ratios can also be predicted from our calculations of the percentage overlap that intervals have with the subtelomeric and pericentromeric regions. For example, the only intervals with an mCO:fCO of less than 2.0 (i.e. CTL1.2; CTL2.4; 3162; Table 2), are those that have no overlap with a subtelomeric region (Table 1), which are known for their concentration of male CO hotspots (Giraut *et al.* 2011).

## Discussion

### Levels of heterochiasmy in Arabidopsis likely show a wide range of intrachromosomal responses to parental age

Our results provide new insights into a previous finding that heterochiasmy in *Arabidopsis* decreases with age (Toyota *et al.* 2011). The current study, only the second to address the topic, showed that in eight out of eight intervals, heterochiasmy did not change with age. However, we do not believe that our results challenge the previous finding. As we will discuss below, we would accept that on a genomic or chromosomal basis, the ratio mCO:fCO is likely to decrease with age; but, our results, when considered together with previous work, suggest that, within any given chromosome, the ratio is unlikely to decrease at many locations. That is, we propose that heterochiasmy in *Arabidopsis* shows a wide spectrum of intrachromosomal responses to age, including, at some locations, no response at all.

For all eight intervals studied here, distributed across all five each chromosomes, neither did mCO:fCO, nor its component rates, change with age. The reliability of our ratio estimations is supported by the good agreement of our results with the rates and ratios that can be calculated using the values of mCO and fCO that Giraut *et al.* (2011) provide for multiple locations within each interval. Some discrepancies might be expected given the different genetic background of the accessions used in the two studies. The relative values of the measured rates and ratios are also as predicted, as accurately as can be expected, by the extent of overlap between an interval and the subtelomeric and pericentromeric regions of the chromosome.

The likelihood that heterochiasmy shows intrachromosomal variation in its response to age is supported by considering our results together with those of Li *et al*. (2017) and Toyota *et al.* (2011). Li *et al*. (2017) studied nine intervals (none of which corresponded to any of our eight) and found that mCO did not change with age in five of nine intervals studied, and increased in two; significantly, there were two cases where, in a single chromosome, mCO changed significantly with age for one interval but not another (Figure 3, Li *et al.* 2017). Although they did not study the response of fCO, the intrachromosomal variation in the response of mCO would also likely lead to variation in the response of mCO:fCO for the intervals studied. Further, in the study of Toyota *et al.* (2011), there was considerable variation, across the chromosomes, in the response of mCO:fCO to age. This was primarily driven by variation in the degree to which fCO increased with age; this increase varied from 1% in chromosome 4 to 14% in chromosome 3 (analysis of Table 5; Toyota *et al.* 2011). Likewise, in the same study, while the response of mCO to age did not change significantly when all five chromosomes were considered together, it did increase by 7% in both chromosomes 1 and 5 (analysis of Table 5; Toyota *et al.* 2011). It is likely that these interchromosomal variations are accompanied by variation at the intrachromosomal level. For example, in chromosome 1, where Toyota *et al.* (2011) found no change in the response of mCO, and only a 1% average change for fCO, there must be many locations where the ratio mCO : fCO did not change with age at all - as we found for all intervals in our study. Also of note here is the study of Francis *et al.* (2007), which found that mCO did not change in response to age.

From our results, together with those of the two previous studies mentioned above (Toyota *et al.* 2011; Li *et al.* 2017), one can propose the following: that as we consider each location along a chromosome of *Arabidopsis* in turn, we will find that: (1) the rates of male and female meiosis will change (as shown by Giraut *et al.* 2011), as will, for many cases, the degree of heterochiasmy; and (2) the sensitivity of rates/ratio to age will change, such that the percentage response to age at any one location will sit somewhere along a broad continuum of values, the starting point of which is zero.

### Which meiotic processes could be responsible for intrachromosomal variability in the response of heterochiasmy to age?

That such a spectrum of age x gender responsiveness exists within a chromosome is not unexpected. A chromosome is highly heterogeneous entity in many respects, and this heterogeneity takes differing forms in male and female meiosis. For example, in humans, synapsis initiation sites are found near the telomeres in male meiosis (Brown *et al.* 2005), but interstitially in females (Lynn *et al.* 2004). Since synapsis initiation sites are also sites for CO (Choi and Henderson 2015), this means that: (a) we can expect different rates of male and female CO at two more or less defined locations within the chromosome; and (b) since each of those locations will vary epigenetically in male and female meiosis (i.e. supporting or not supporting synapsis initiation), this opens up the possibility that each will also respond differently to the many cellular changes that accompany ageing.

In *Arabdiopsis*, there is not the same tight coupling between synapsis initiation and CO (Chelysheva *et al.* 2007). However, we can be certain that, if we compare any pair of corresponding locations in a chromosome undergoing male or female meiosis, there is a high probability that their epigenetic environments will differ. A visual scan of the male and female recombination landscapes of any of the five chromosomes (Figure 4, Giraut *et al.* 2011) shows that the rates, and the ratio, mCO: CO, vary frequently and dramatically from location to location. Their analysis found that amongst 380 locations in the genome, more than half were significantly hot or cold, in terms of male or female CO rates. The hotspots are thus necessarily not just concentrated in one area of the chromosome: e.g. 27/40 of the male hotspots occur away from the telomeres (Figure 4, Giraut *et al.* 2011). Given that the chromosomes involved in male and female meiosis have the same sequence, a mechanistic explanation for the difference between the CO rates at the same location must lie in their exposure to different epigenetic conditions. As with the human example described in the previous paragraph, this will create the potential for intrachromosomal and sex-based variation in the responses to age.

The mechanisms that might lead to these local chromosomal differences in mCO and fCO, or the cumulative differences they bring about at the genome level, have been the focus of many studies in *Arabidopsis*, and these have indicated a range of epigenetic mechanisms that could provide a ‘substrate’ for some or all of the intrachromosomal variation in age x gender responses. What is interesting to consider, for our purposes, is which of the mechanisms might be both: (1) significantly important for the differences in male and female CO rates; and (2) highly responsive to age.

Perhaps the strongest candidate is the degree of chromatin compaction, including -for each ‘level’ of compaction-the associated molecular players that maintain that level. In *Arabidopsis*, female chromosomes are markedly more compact, as indexed by their much shorter synaptonemal complexes (Drouaud *et al.* 2007); synaptonemal complex length is a known indicator of chromatin compaction and a predictor of meiotic combination rate in *Arabidopsis*, and other species (Kleckner 2006; Drouaud *et al.* 2007; Brachet *et al.* 2012; Zickler and Kleckner 2015; Wang *et al.* 2016; Modliszewski and Copenhaver 2017). Modelling studies have also indicated that sex variations in chromosomal structural axis length (which is related to synaptonemal complex length) are sufficient to explain the sex variation in CO rates in *Arabidopsis* (Zickler and Kleckner 2015). Control of chromatin organisation ensures that chromosomes have the right length, and other structural features, that are critical to the proper alignment of daughter chromatids, and thus crossing over itself (Brachet *et al.* 2012; Stapley *et al.* 2017).

Chromatin restructuring, particularly of heterochromatin, is also well established as one of the common features of cellular ageing and senescence in animals; chromatin becomes locally more or less compact with age (Vaquero *et al.* 2003; Swanson *et al.* 2015). In plants, less is known, but methylation patterns, which are associated with both chromatin compaction and CO rates, do change with age: in *Arabidopsis*, for example, ageing is accompanied by DNA demethylation (Ogneva *et al.* 2016); also, in *Arabidopsis* leaf senescence, heterochromatin disintegrates (Ay *et al.* 2009). Another feature of chromatin compaction that makes it an attractive candidate is that the level of compaction naturally varies along the length of the chromosome, being negatively correlated with gene density (Brachet *et al.* 2012). This, together, with the known global differences between compaction levels in male and female male meiotic cells, suggests that local variations in compaction level could provide the rich epigenetic substrate needed for the extensive age x gender intrachromosomal variation indicated by the current and previous findings.

## Conclusions

Our results help, we believe, to clarify how the levels of heterochiasmy in *Arabidopsis* respond to age. From previous work, it might be concluded that the level of disparity between rates will drop universally across the genome, but we suggest that the magnitude, perhaps even the direction, of the response depends on which part of the genome is sampled. This has implications for other studies that look at the interactive effects of multiple factors on any given meiotic response, indicating the possibility of different findings if global or local sampling approaches are adopted. Our results will also be of interest to researchers who are looking for ways to bring about increased CO rates in specific regions of the genome (Fernandes *et al.* 2018). If any given approach to inducing a rate increase has been rejected in the past because it did not lead to a global rate increase, the possibility nevertheless remains that the approach might induce increased CO rates in a local genomic region of interest.

## Supporting information

GSA

GSA

## Author Contributions

Conceived and designed the experiments: RS, AKS and RB. Performed the experiments and compiled the data: RS. Analysed the data: RS, AKS, GJH, and RB. Wrote the paper: RS, AKS, GJH and RB.

## Acknowledgements

We would like to thank Avraham A. Levy (Weizmann Institute of Science, Israel) for providing seeds of MR detector lines. We would like to thank the Arabidopsis Biological Resource Center (The Ohio State University, USA) for providing traffic line seeds.

## Literature Cited

Alves I., A. A. Houle, J. G. Hussin, and P. Awadalla, 2017 The impact of recombination on human mutation load and disease. Philos. Trans. R. Soc. B Biol. Sci. 372: 20160465. https://doi.org/10.1098/rstb.2016.0465

Ay N., K. Irmler, A. Fischer, R. Uhlemann, G. Reuter, et al., 2009 Epigenetic programming via histone methylation at WRKY53 controls leaf senescence in Arabidopsis thaliana. Plant J. 58: 333–346. https://doi.org/10.1111/j.0960-7412.2009.03782.x

Brachet E., V. Sommermeyer, and V. Borde, 2012 Interplay between modifications of chromatin and meiotic recombination hotspots. Biol. Cell 104: 51–69. https://doi.org/10.1111/boc.201100113

Bretz F., T. Hothorn, and P. Westfall, 2016 Multiple comparisons using R. Chapman and Hall/CRC.

Brown P. W., L. Judis, E. R. Chan, S. Schwartz, A. Seftel, et al., 2005 Meiotic synapsis proceeds from a limited number of subtelomeric sites in the human male. Am. J. Hum. Genet. 77: 556–566.

Charlesworth D., and X. Vekemans, 2005 How and when did Arabidopsis thaliana become highly self-fertilising. BioEssays 27: 472–476. https://doi.org/10.1002/bies.20231

Chelysheva L., G. Gendrot, D. Vezon, M.-P. Doutriaux, R. Mercier, et al., 2007 Zip4/Spo22 is required for Class I CO formation but not for synapsis completion in Arabidopsis thaliana. PLOS Genet. 3: e83. https://doi.org/10.1371/journal.pgen.0030083

Chiang T., R. M. Schultz, and M. A. Lampson, 2012 Meiotic origins of maternal age-related aneuploidy1. Biol. Reprod. 86. https://doi.org/10.1095/biolreprod.111.094367

Choi K., and I. R. Henderson, 2015 Meiotic recombination hotspots – a comparative view. Plant J. 83: 52–61. https://doi.org/10.1111/tpj.12870

Drouaud J., V. Zanni, and D. Brunel, 2007 Sex-specific crossover distributions and variations in interference level along Arabidopsis thaliana chromosome 4. PLoS Genet. 3: 12.

Fernandes J. B., M. Séguéla-Arnaud, C. Larchevêque, A. H. Lloyd, and R. Mercier, 2018 Unleashing meiotic crossovers in hybrid plants. Proc. Natl. Acad. Sci. 115: 2431. https://doi.org/10.1073/pnas.1713078114

Francis K. E., S. Y. Lam, B. D. Harrison, A. L. Bey, L. E. Berchowitz, et al., 2007 Pollen tetrad-based visual assay for meiotic recombination in Arabidopsis. Proc. Natl. Acad. Sci. 104: 3913–3918. https://doi.org/10.1073/pnas.0608936104

Gabriel K. R., 1969 Simultaneous test procedures--some theory of multiple comparisons. Ann. Math. Stat. 224–250.

Giraut L., M. Falque, J. Drouaud, L. Pereira, O. C. Martin, et al., 2011 Genome-wide crossover distribution in Arabidopsis thaliana meiosis reveals sex-specific patterns along chromosomes, (M. Lichten, Ed.). PLoS Genet. 7: e1002354. https://doi.org/10.1371/journal.pgen.1002354

Halldorsson B. V., M. T. Hardarson, B. Kehr, U. Styrkarsdottir, A. Gylfason, et al., 2016 The rate of meiotic gene conversion varies by sex and age. Nat. Genet. 48: 1377–1384. https://doi.org/10.1038/ng.3669

Hayman D. L., and P. A. Parsons, 1962 The effect of temperature, age and an inversion on recombination values and interference in the X-chromosome of Drosophila melanogaster. Genetica 32: 74–88. https://doi.org/10.1007/BF01816087

Hussin J., M.-H. Roy-Gagnon, R. Gendron, G. Andelfinger, and P. Awadalla, 2011 Age-dependent recombination rates in human pedigrees, (G. McVean, Ed.). PLoS Genet. 7: e1002251. https://doi.org/10.1371/journal.pgen.1002251

John A., K. Vinayan, and J. Varghese, 2016 Achiasmy: Male fruit flies are not ready to mix. Front. Cell Dev. Biol. 4. https://doi.org/10.3389/fcell.2016.00075

Kleckner N., 2006 Chiasma formation: chromatin/axis interplay and the role(s) of the synaptonemal complex. Chromosoma 115: 175.

Lenormand T., 2003 The evolution of sex dimorphism in recombination. Genetics 163: 811–822.

Lenormand T., and J. Dutheil, 2005 Recombination Difference between sexes: A role for haploid selection. PLoS Biol. 3: e63. https://doi.org/10.1371/journal.pbio.0030063

Li F., N. De Storme, and D. Geelen, 2017 Dynamics of male meiotic recombination frequency during plant development using Fluorescent Tagged Lines in Arabidopsis thaliana. Sci. Rep. 7: 42535. https://doi.org/10.1038/srep42535

López E., M. Pradillo, C. Oliver, C. Romero, N. Cuñado, et al., 2012 Looking for natural variation in chiasma frequency in Arabidopsis thaliana. J. Exp. Bot. 63: 887–894. https://doi.org/10.1093/jxb/err319

Lynn A., T. Ashley, and T. Hassold, 2004 Variation in human meiotic recombination. Annu Rev Genomics Hum Genet 5: 317–349.

Martin H. C., R. Christ, J. G. Hussin, J. O’Connell, S. Gordon, et al., 2015 Multicohort analysis of the maternal age effect on recombination. Nat. Commun. 6: 7846. https://doi.org/10.1038/ncomms8846

McKee B. D., R. Yan, and J.-H. Tsai, 2012 Meiosis in male Drosophila. Spermatogenesis 2: 167–184. https://doi.org/10.4161/spmg.21800

Melamed-Bessudo C., E. Yehuda, A. R. Stuitje, and A. A. Levy, 2005 A new seed-based assay for meiotic recombination in Arabidopsis thaliana: Meiotic recombination in Arabidopsis thaliana. Plant J. 43: 458–466. https://doi.org/10.1111/j.1365-313X.2005.02466.x

Modliszewski J. L., and G. P. Copenhaver, 2017 Meiotic recombination gets stressed out: CO frequency is plastic under pressure. Curr. Opin. Plant Biol. 36: 95–102. https://doi.org/10.1016/j.pbi.2016.11.019

Nagaoka S. I., T. J. Hassold, and P. A. Hunt, 2012 Human aneuploidy: mechanisms and new insights into an age-old problem. Nat. Rev. Genet. Lond. 13: 493–504. http://dx.doi.org.ezproxy.uws.edu.au/10.1038/nrg3245

Nelder J. A., and R. W. Wedderburn, 1972 Generalized linear models. J. R. Stat. Soc. Ser. Gen. 135: 370–384.

Ogneva Z. V., A. S. Dubrovina, and K. V. Kiselev, 2016 Age-associated alterations in DNA methylation and expression of methyltransferase and demethylase genes in Arabidopsis thaliana. Biol. Plant. 60: 628–634. https://doi.org/10.1007/s10535-016-0638-y

Ritz K. R., M. A. F. Noor, and N. D. Singh, 2017 Variation in recombination rate: Adaptive or not? Trends Genet. 33: 364–374. https://doi.org/10.1016/j.tig.2017.03.003

Saini R., A. K. Singh, S. Dhanapal, T. H. Saeed, G. J. Hyde, et al., 2017 Brief temperature stress during reproductive stages alters meiotic recombination and somatic mutation rates in the progeny of Arabidopsis. BMC Plant Biol. 17: 103. https://doi.org/10.1186/s12870-017-1051-1

Satomura K., N. Osada, and T. Endo, 2019 Achiasmy and sex chromosome evolution. Ecol. Genet. Genomics 13: 100046. https://doi.org/10.1016/j.egg.2019.100046

Singh A. K., T. Bashir, C. Sailer, V. Gurumoorthy, A. M. Ramakrishnan, et al., 2015 Parental age affects somatic mutation rates in the progeny of flowering plants. Plant Physiol. 168: 247–257. https://doi.org/10.1104/pp.15.00291

Stapley J., P. G. D. Feulner, S. E. Johnston, A. W. Santure, and C. M. Smadja, 2017 Variation in recombination frequency and distribution across eukaryotes: patterns and processes. Philos. Trans. R. Soc. B Biol. Sci. 372: 20160455. https://doi.org/10.1098/rstb.2016.0455

Swanson E. C., L. M. Rapkin, D. P. Bazett-Jones, and J. B. Lawrence, 2015 Unfolding the story of chromatin organization in senescent cells. Nucleus 6: 254–260. https://doi.org/10.1080/19491034.2015.1057670

Toyota M., K. Matsuda, T. Kakutani, M. Terao Morita, and M. Tasaka, 2011 Developmental changes in crossover frequency in Arabidopsis: Sex and age dependence of crossover frequency. Plant J. 65: 589–599. https://doi.org/10.1111/j.1365-313X.2010.04440.x

Vaquero A., A. Loyola, and D. Reinberg, 2003 The constantly changing face of chromatin. Sci. Aging Knowl. Environ. 2003: 4.

Wang Z., B. Shen, J. Jiang, J. Li, and L. Ma, 2016 Effect of sex, age and genetics on crossover interference in cattle. Sci. Rep. 6: 37698. https://doi.org/10.1038/srep37698

Wu G., G. Rossidivito, T. Hu, Y. Berlyand, and R. S. Poethig, 2015 Traffic lines: new tools for genetic analysis in Arabidopsis thaliana. Genetics 200: 35–45.

Zickler D., and N. Kleckner, 2015 Recombination, pairing, and synapsis of homologs during meiosis. Cold Spring Harb. Perspect. Biol. 7. https://doi.org/10.1101/cshperspect.a016626

