## Supplementary material for "Sex-specific crossover rates did not change with parental age in *Arabidopsis*": GSA

**Table S1: CO rates in the progeny of a cross between parents of CO detector line and wild type Columbia plants of different ages (CTL1.2)**

| **Cross** | **CO Rate (R)/Seeds per replicate (S)** | | | | **Mean CO Rate** | |
| --- | --- | --- | --- | --- | --- | --- |
|  | R1/S1 | R2/S2 | R3/S3 | |  |  |
| CTL1.2 40 X Col 40 | 31.29/930 | 30.49/469 | 25.14/366 | | 28.9733 | |
| Col 40 X CTL1.2 40 | 29.02/485 | 28.93/473 | 26.4/590 | | 28.1167 | |
| CTL1.2 45 X Col 45 | 25.19/1137 | 25.52/674 | 25.36/606 | | 25.3567 | |
| Col 45 X CTL1.2 45 | 31.22/721 | 30.89/690 | 29.55/175 | | 30.5533 | |
| CTL1.2 50 X Col 50 | 33.81/258 | 31.5/286 | 26.45/321 | | 30.5867 | |
| Col 50 X CTL1.2 50 | 39.11/363 | 37.82/320 | 37.14/432 | | 38.0233 | |
| CTL1.2 55 X Col 55 | 34.71/1214 | 29.38/913 | 27.77/423 | | 30.62 | |
| Col 55 X CTL1.2 55 | 36.42/486 | 34.1/541 | 33.14/398 | | 34.5533 | |
| **Comparison between different groups** | | | | | | |
| Cross | | | | P Value | | Significance |
| CTL1.2 40 X Col 40-CTL1.2 45 X Col 45 | | | | 0.9997 | | NS |
| CTL1.2 40 X Col 40-CTL1.2 50 X Col 50 | | | | 0.4889 | | NS |
| CTL1.2 40 X Col 40-CTL1.2 55 X Col 55 | | | | 0.9874 | | NS |
| CTL1.2 45 X Col 45-CTL1.2 50 X Col 50 | | | | 0.7912 | | NS |
| CTL1.2 45 X Col 45-CTL1.2 55 X Col 55 | | | | 0.8774 | | NS |
| CTL1.2 50 X Col 50-CTL1.2 55 X Col 55 | | | | 0.0911 | | NS |
| Col 40 X CTL1.2 40- Col 45 X CTL1.2 45 | | | | 0.0610 | | NS |
| Col 40 X CTL1.2 40- Col 50 X CTL1.2 50 | | | | 1.0000 | | NS |
| Col 40 X CTL1.2 40- Col 55 X CTL1.2 55 | | | | 0.3682 | | NS |
| Col 45 X CTL1.2 45- Col 50 X CTL1.2 50 | | | | 0.5130 | | NS |
| Col 45 X CTL1.2 45- Col 55 X CTL1.2 55 | | | | 0.5435 | | NS |
| Col 50 X CTL1.2 50- Col 55 X CTL1.2 55 | | | | 0.3787 | | NS |
| CTL1.2 40 X Col 40- Col 40 X CTL1.2 40 | | | | 0.9858 | | NS |
| CTL1.2 45 X Col 45- Col 45 X CTL1.2 45 | | | | <0.01 | | ** |
| CTL1.2 50 X Col 50- Col 50 X CTL1.2 50 | | | | 0.0837 | | NS |
| CTL1.2 55 X Col 55- Col 55 X CTL1.2 55 | | | | 0.3576 | | NS |

**Table S2: CO rates in the progeny of a cross between parents of CO detector line and wild type Columbia plants of different ages (CTL1.18)**

| **Cross** | **CO Rate (R)/Seeds per replicate (S)** | | | | **Mean CO Rate** | |
| --- | --- | --- | --- | --- | --- | --- |
|  | R1/S1 | R2/S2 | R3/S3 | |  |  |
| CTL1.18 40 X Col 40 | 6.4/750 | 8.03/498 | 9.6/957 | | 8.01 | |
| Col 40 X CTL1.18 40 | 5.71/479 | 6.85/672 | 5.24/693 | | 5.93 | |
| CTL1.18 45 X Col 45 | 8.22/840 | 10.16/1008 | 9.19/782 | | 9.19 | |
| Col 45 X CTL1.18 45 | 6.84/604 | 9.75/906 | 7.76/509 | | 8.12 | |
| CTL1.18 50 X Col 50 | 18.58/365 | 18.75/246 | 19.49/380 | | 18.94 | |
| Col 50 X CTL1.18 50 | 20.03/423 | 15.67/433 | 15.72/387 | | 17.14 | |
| CTL1.18 55 X Col 55 | 18.44/921 | 16.42/636 | 17.43/323 | | 17.43 | |
| Col 55 X CTL1.18 55 | 18.73/299 | 17.87/431 | 18.3/342 | | 18.3 | |
| **Comparison between different groups** | | | | | | |
| Cross | | | | P Value | | Significance |
| CTL1.18 40 X Col 40-CTL1.18 45 X Col 45 | | | | 0.4502 | | NS |
| CTL1.18 40 X Col 40-CTL1.18 50 X Col 50 | | | | 0.9375 | | NS |
| CTL1.18 40 X Col 40-CTL1.18 55 X Col 55 | | | | 1.0000 | | NS |
| CTL1.18 45 X Col 45-CTL1.18 50 X Col 50 | | | | 0.0556 | | NS |
| CTL1.18 45 X Col 45-CTL1.18 55 X Col 55 | | | | 0.3839 | | NS |
| CTL1.18 50 X Col 50-CTL1.18 55 X Col 55 | | | | 0.9618 | | NS |
| Col 40 X CTL1.18 40- Col 45 X CTL1.18 45 | | | | 0.6316 | | NS |
| Col 40 X CTL1.18 40- Col 50 X CTL1.18 50 | | | | 0.8060 | | NS |
| Col 40 X CTL1.18 40- Col 55 X CTL1.18 55 | | | | 0.9982 | | NS |
| Col 45 X CTL1.18 45- Col 50 X CTL1.18 50 | | | | 1.0000 | | NS |
| Col 45 X CTL1.18 45- Col 55 X CTL1.18 55 | | | | 0.9427 | | NS |
| Col 50 X CTL1.18 50- Col 55 X CTL1.18 55 | | | | 0.9883 | | NS |
| CTL1.18 40 X Col 40- Col 40 X CTL1.18 40 | | | | <0.001 | | *** |
| CTL1.18 45 X Col 45- Col 45 X CTL1.18 45 | | | | <0.001 | | *** |
| CTL1.18 50 X Col 50- Col 50 X CTL1.18 50 | | | | <0.001 | | *** |
| CTL1.18 55 X Col 55- Col 55 X CTL1.18 55 | | | | <0.001 | | *** |

**Table S3: CO rates in the progeny of a cross between parents of CO detector line and wild type Columbia plants of different ages (CTL2.4)**

| **Cross** | **CO Rate (R)/Seeds per replicate (S)** | | | | **Mean CO Rate** | |
| --- | --- | --- | --- | --- | --- | --- |
|  | R1/S1 | R2/S2 | R3/S3 | |  |  |
| CTL2.4 40 X Col 40 | 12.48/858 | 15.67/351 | 11.74/281 | | 13.2967 | |
| Col 40 X CTL2.4 40 | 14.69/602 | 16.43/127 | 12.99/155 | | 14.7033 | |
| CTL2.4 45 X Col 45 | 10.94/708 | 13.18/347 | 10.42/308 | | 11.5133 | |
| Col 45 X CTL2.4 45 | 14.76/851 | 13.58/595 | 14.17/345 | | 14.17 | |
| CTL2.4 50 X Col 50 | 15.61/594 | 20.47/645 | 23.88/499 | | 19.9867 | |
| Col 50 X CTL2.4 50 | 15.16/555 | 16.3/205 | 17.1/421 | | 16.1867 | |
| CTL2.4 55 X Col 55 | 16.04/271 | 19.51/302 | 17.78/345 | | 17.7767 | |
| Col 55 X CTL2.4 55 | 16.76/105 | 17.5/203 | 18.1/178 | | 17.4533 | |
| **Comparison between different groups** | | | | | | |
| Cross | | | | P Value | | Significance |
| CTL2.4 40 X Col 40-CTL2.4 45 X Col 45 | | | | 0.982 | | NS |
| CTL2.4 40 X Col 40-CTL2.4 50 X Col 50 | | | | 0.934 | | NS |
| CTL2.4 40 X Col 40-CTL2.4 55 X Col 55 | | | | 0.999 | | NS |
| CTL2.4 45 X Col 45-CTL2.4 50 X Col 50 | | | | 0.414 | | NS |
| CTL2.4 45 X Col 45-CTL2.4 55 X Col 55 | | | | 1.000 | | NS |
| CTL2.4 50 X Col 50-CTL2.4 55 X Col 55 | | | | 0.647 | | NS |
| Col 40 X CTL2.4 40- Col 45 X CTL2.4 45 | | | | 0.202 | | NS |
| Col 40 X CTL2.4 40- Col 50 X CTL2.4 50 | | | | 0.822 | | NS |
| Col 40 X CTL2.4 40- Col 55 X CTL2.4 55 | | | | 0.700 | | NS |
| Col 45 X CTL2.4 45- Col 50 X CTL2.4 50 | | | | 0.964 | | NS |
| Col 45 X CTL2.4 45- Col 55 X CTL2.4 55 | | | | 0.990 | | NS |
| Col 50 X CTL2.4 50- Col 55 X CTL2.4 55 | | | | 1.000 | | NS |
| CTL2.4 40 X Col 40- Col 40 X CTL2.4 40 | | | | <0.01 | | *** |
| CTL2.4 45 X Col 45- Col 45 X CTL2.4 45 | | | | 0.975 | | NS |
| CTL2.4 50 X Col 50- Col 50 X CTL2.4 50 | | | | <0.01 | | ** |
| CTL2.4 55 X Col 55- Col 55 X CTL2.4 55 | | | | 0.376 | | NS |

**Table S4: CO rates in the progeny of a cross between parents of CO detector line and wild type Columbia plants of different ages (CTL4.7)**

| **Cross** | **CO Rate (R)/Seeds per replicate (S)** | | | | **Mean CO Rate** | |
| --- | --- | --- | --- | --- | --- | --- |
|  | R1/S1 | R2/S2 | R3/S3 | |  |  |
| CTL4.7 40 X Col 40 | 9.9/777 | 7.75/619 | 9.3/570 | | 8.9833 | |
| Col 40 X CTL4.7 40 | 21.24/339 | 26.59/267 | 23.91/304 | | 23.9133 | |
| CTL4.7 45 X Col 45 | 7.43/579 | 8.83/668 | 9.12/1162 | | 8.46 | |
| Col 45 X CTL4.7 45 | 24.43/700 | 23.76/884 | 25.36/698 | | 24.5167 | |
| CTL4.7 50 X Col 50 | 8.53/598 | 8.16/457 | 8.34/387 | | 8.3433 | |
| Col 50 X CTL4.7 50 | 23.28/421 | 22.77/505 | 22.89/380 | | 22.98 | |
| CTL4.7 55 X Col 55 | 8.47/956 | 9.34/996 | 8.9/742 | | 8.9033 | |
| Col 55 X CTL4.7 55 | 20.45/220 | 21.23/345 | 22.15/412 | | 21.2767 | |
| **Comparison between different groups** | | | | | | |
| Cross | | | | P Value | | Significance |
| CTL4.7 40 X Col 40-CTL4.7 45 X Col 45 | | | | 0.999 | | NS |
| CTL4.7 40 X Col 40-CTL4.7 50 X Col 50 | | | | 0.996 | | NS |
| CTL4.7 40 X Col 40-CTL4.7 55 X Col 55 | | | | 1.000 | | NS |
| CTL4.7 45 X Col 45-CTL4.7 50 X Col 50 | | | | 1.000 | | NS |
| CTL4.7 45 X Col 45-CTL4.7 55 X Col 55 | | | | 0.999 | | NS |
| CTL4.7 50 X Col 50-CTL4.7 55 X Col 55 | | | | 0.998 | | NS |
| Col 40 X CTL4.7 40- Col 45 X CTL4.7 45 | | | | 0.997 | | NS |
| Col 40 X CTL4.7 40- Col 50 X CTL4.7 50 | | | | 0.964 | | NS |
| Col 40 X CTL4.7 40- Col 55 X CTL4.7 55 | | | | 0.070 | | NS |
| Col 45 X CTL4.7 45- Col 50 X CTL4.7 50 | | | | 0.664 | | NS |
| Col 45 X CTL4.7 45- Col 55 X CTL4.7 55 | | | | 0.009 | | NS |
| Col 50 X CTL4.7 50- Col 55 X CTL4.7 55 | | | | 0.539 | | NS |
| CTL4.7 40 X Col 40- Col 40 X CTL4.7 40 | | | | <0.001 | | *** |
| CTL4.7 45 X Col 45- Col 45 X CTL4.7 45 | | | | <0.001 | | *** |
| CTL4.7 50 X Col 50- Col 50 X CTL4.7 50 | | | | <0.001 | | *** |
| CTL4.7 55 X Col 55- Col 55 X CTL4.7 55 | | | | <0.001 | | *** |

**Table S5: CO rates in the progeny of a cross between parents of CO detector line and wild type Columbia plants of different ages (Col3-4/20)**

| **Cross** | **CO Rate (R)/Seeds per replicate (S)** | | | | | | | | | **Mean CO Rate** |
| --- | --- | --- | --- | --- | --- | --- | --- | --- | --- | --- |
|  | R1/S1 | R2/S2 | R3/S3 | R4/S4 | R5/S5 | R6/S6 | R7/S7 | R8/S8 | R9/S9 |  |
| Col3-4/20 40 X Col 40 | 8.07 | 8.61 | 10 | 7.46 | 8.95 | 12 |  |  |  | 9.18167 |
|  | 233 | 511 | 320 | 268 | 268 | 475 |  |  |  |  |
| Col 40 X Col3-4/20 40 | 23.29 | 20.72 | 21.78 | 27.51 | 21.9 | 23.67 |  |  |  | 23.145 |
|  | 176 | 193 | 303 | 378 | 242 | 511 |  |  |  |  |
| Col3-4/20 45 X Col 45 | 8.82 | 8.4 | 8.01 | 10.26 | 11.74 | 9.57 | 8.36 | 6.61 | 8.19 | 8.97125 |
|  | 612 | 702 | 287 | 565 | 681 | 355 | 622 | 590 | 122 |  |
| Col 45 X Col3-4/20 45 | 23.43 | 20.45 | 20.66 | 25.36 | 24.14 | 25 | 28.01 | 27.07 | 27.54 | 24.265 |
|  | 431 | 396 | 150 | 138 | 758 | 268 | 257 | 314 | 511 |  |
| Col3-4/20 50 X Col 50 | 9.57 | 7.4 | 7.27 | 10.97 | 9.85 | 9.64 |  |  |  | 9.44 |
|  | 658 | 378 | 522 | 401 | 406 | 311 |  |  |  |  |
| Col 50 X Col3-4/20 50 | 26.06 | 27.19 | 22.49 | 20.68 | 28.44 | 23.61 |  |  |  | 24.6988 |
|  | 211 | 1140 | 609 | 290 | 334 | 199 |  |  |  |  |
| Col3-4/20 55 X Col 55 | 9.6 | 11.48 | 5.75 | 6.26 | 6.8 | 7.35 | 8.87 | 8.04 | 8.53 | 8.01875 |
|  | 500 | 444 | 191 | 527 | 354 | 340 | 248 | 261 | 668 |  |
| Col 55 X Col3-4/20 55 | 20.27 | 30.43 | 18.91 | 21.5 | 24 | 22.05 | 23.39 | 32.03 | 27.78 | 24.0725 |
|  | 148 | 138 | 185 | 413 | 229 | 217 | 218 | 231 | 511 |  |
| **Comparison between different groups** | | | | | | | | | | |
| Cross | | | | | | P Value | | Significance | | |
| Col3-4/20 40 X Col 40-Col3-4/20 45 X Col 45 | | | | | | 1.000 | | Non-Significant (NS) | | |
| Col3-4/20 40 X Col 40-Col3-4/20 50 X Col 50 | | | | | | 1.000 | | NS | | |
| Col3-4/20 40 X Col 40-Col3-4/20 55 X Col 55 | | | | | | 0.996 | | NS | | |
| Col3-4/20 45 X Col 45-Col3-4/20 50 X Col 50 | | | | | | 1.000 | | NS | | |
| Col3-4/20 45 X Col 45-Col3-4/20 55 X Col 55 | | | | | | 0.999 | | NS | | |
| Col3-4/20 50 X Col 50-Col3-4/20 55 X Col 55 | | | | | | 0.970 | | NS | | |
| Col 40 X Col3-4/20 40- Col 45 X Col3-4/20 45 | | | | | | 0.976 | | NS | | |
| Col 40 X Col3-4/20 40- Col 50 X Col3-4/20 50 | | | | | | 0.981 | | NS | | |
| Col 40 X Col3-4/20 40- Col 55 X Col3-4/20 55 | | | | | | 1.000 | | NS | | |
| Col 45 X Col3-4/20 45- Col 50 X Col3-4/20 50 | | | | | | 1.000 | | NS | | |
| Col 45 X Col3-4/20 45- Col 55 X Col3-4/20 55 | | | | | | 0.712 | | NS | | |
| Col 50 X Col3-4/20 50- Col 55 X Col3-4/20 55 | | | | | | 0.735 | | NS | | |
| Col3-4/20 40 X Col 40- Col 40 X Col3-4/20 40 | | | | | | <0.001 | | *** | | |
| Col3-4/20 45 X Col 45- Col 45 X Col3-4/20 45 | | | | | | <0.001 | | *** | | |
| Col3-4/20 50 X Col 50- Col 50 X Col3-4/20 50 | | | | | | <0.001 | | *** | | |
| Col3-4/20 55 X Col 55- Col 55 X Col3-4/20 55 | | | | | | <0.001 | | *** | | |

**Table S6: CO rates in the progeny of a cross between parents of CO detector line and wild type Columbia plants of different ages (3158)**

| **Cross** | **CO Rate (R)/Seeds per replicate (S)** | | | | | | | | | **Mean CO Rate** |
| --- | --- | --- | --- | --- | --- | --- | --- | --- | --- | --- |
|  | R1/S1 | R2/S2 | R3/S3 | R4/S4 | R5/S5 | R6/S6 | R7/S7 | R8/S8 | R9/S9 |  |
| 3158 40 X Col 40 | 10 | 9.53 | 15.41 | 9.32 | 11.77 | 14.47 |  |  |  | 11.75 |
|  | 580 | 619 | 597 | 504 | 781 | 995 |  |  |  |  |
| Col 40 X 3158 40 | 25.21 | 26.33 | 26.33 | 29.54 | 26.99 | 30.16 |  |  |  | 27.4266 |
|  | 230 | 281 | 346 | 264 | 363 | 885 |  |  |  |  |
| 3158 45 X Col 45 | 12 | 9.94 | 7.98 | 9.51 | 10.99 | 10.2 | 12.75 | 12.58 | 9.78 | 10.6366 |
|  | 500 | 784 | 213 | 389 | 646 | 147 | 1215 | 858 | 276 |  |
| Col 45 X 3158 45 | 23.81 | 24.6 | 25.62 | 25.24 | 26.29 | 27.89 | 22.89 | 24.35 | 23.62 | 24.9233 |
|  | 953 | 263 | 111 | 103 | 559 | 294 | 83 | 386 | 452 |  |
| 3158 50 X Col 50 | 9.1 | 10.59 | 10.57 | 8.27 | 12.88 | 11.23 |  |  |  | 10.5577 |
|  | 670 | 151 | 709 | 810 | 326 | 463 |  |  |  |  |
| Col 50 X 3158 50 | 25.43 | 22.92 | 25.18 | 25.85 | 26.13 | 25.99 |  |  |  | 25.4966 |
|  | 287 | 253 | 1100 | 294 | 528 | 245 |  |  |  |  |
| 3158 55 X Col 55 | 12.83 | 14.67 | 9.9 | 5.98 | 12.92 | 10.26 | 8.69 | 12.19 | 7.9 | 10.5933 |
|  | 1067 | 1097 | 323 | 167 | 565 | 224 | 276 | 1009 | 329 |  |
| Col 55 X 3158 55 | 28.12 | 28.42 | 27.82 | 23.69 | 23.69 | 23.69 | 22.3 | 25 | 22.09 | 24.9633 |
|  | 129 | 570 | 345 | 595 | 465 | 130 | 712 | 267 | 352 |  |
| **Comparison between different groups** | | | | | | | | | | |
| Cross | | | | | | P Value | | Significance | | |
| 3158 40 X Col 40-3158 45 X Col 45 | | | | | | 0.932 | | NS | | |
| 3158 40 X Col 40-3158 50 X Col 50 | | | | | | 0.932 | | NS | | |
| 3158 40 X Col 40-3158 55 X Col 55 | | | | | | 0.941 | | NS | | |
| 3158 45 X Col 45-3158 50 X Col 50 | | | | | | 1.000 | | NS | | |
| 3158 45 X Col 45-3158 55 X Col 55 | | | | | | 1.000 | | NS | | |
| 3158 50 X Col 50-3158 55 X Col 55 | | | | | | 1.000 | | NS | | |
| Col 40 X 3158 40- Col 45 X 3158 45 | | | | | | 0.253 | | NS | | |
| Col 40 X 3158 40- Col 50 X 3158 50 | | | | | | 0.598 | | NS | | |
| Col 40 X 3158 40- Col 55 X 3158 55 | | | | | | 0.236 | | NS | | |
| Col 45 X 3158 45- Col 50 X 3158 50 | | | | | | 0.998 | | NS | | |
| Col 45 X 3158 45- Col 55 X 3158 55 | | | | | | 1.000 | | NS | | |
| Col 50 X 3158 50- Col 55 X 3158 55 | | | | | | 0.997 | | NS | | |
| 3158 40 X Col 40- Col 40 X 3158 40 | | | | | | <0.01 | | *** | | |
| 3158 45 X Col 45- Col 45 X 3158 45 | | | | | | <0.01 | | *** | | |
| 3158 50 X Col 50- Col 50 X 3158 50 | | | | | | <0.01 | | *** | | |
| 3158 55 X Col 55- Col 55 X 3158 55 | | | | | | <0.01 | | *** | | |

**Table S7: CO rates in the progeny of a cross between parents of CO detector line and wild type Columbia plants of different ages (3162)**

| **Cross** | **CO Rate (R)/Seeds per replicate (S)** | | | | | | | | | **Mean CO Rate** |
| --- | --- | --- | --- | --- | --- | --- | --- | --- | --- | --- |
|  | R1/S1 | R2/S2 | R3/S3 | R4/S4 | R5/S5 | R6/S6 | R7/S7 | R8/S8 | R9/S9 |  |
| 3162 40 X Col 40 | 14.51 | 19.72 | 17.11 | 17.11 | 16.96 | 16.02 |  |  |  | 16.905 |
|  | 503 | 507 | 495 | 333 | 825 | 699 |  |  |  |  |
| Col 40 X 3162 40 | 14.83 | 22.38 | 18.35 | 23.14 | 16.41 | 20 |  |  |  | 19.185 |
|  | 236 | 210 | 409 | 363 | 262 | 569 |  |  |  |  |
| 3162 45 X Col 45 | 16.41 | 15.09 | 16.18 | 14.25 | 15.57 | 13.51 | 13.11 | 15.45 | 14.46 | 14.8922 |
|  | 798 | 669 | 241 | 505 | 687 | 148 | 900 | 893 | 235 |  |
| Col 45 X 3162 45 | 19.64 | 18.04 | 19.65 | 22.18 | 18.56 | 20.37 | 19.2 | 21.01 | 20.1 | 19.8611 |
|  | 285 | 388 | 234 | 604 | 528 | 354 | 276 | 295 | 284 |  |
| 3162 50 X Col 50 | 16.3 | 10.85 | 14.34 | 11.33 | 11.53 | 13.97 |  |  |  | 12.7944 |
|  | 368 | 258 | 669 | 247 | 390 | 415 |  |  |  |  |
| Col 50 X 3162 50 | 19.17 | 20.51 | 22.87 | 17.51 | 20.09 | 18.8 |  |  |  | 19.4833 |
|  | 219 | 351 | 529 | 394 | 403 | 439 |  |  |  |  |
| 3162 55 X Col 55 | 14.85 | 17.71 | 16.28 | 12.63 | 13.9 | 14.04 | 11.79 | 14.81 | 13.3 | 14.3677 |
|  | 175 | 1287 | 463 | 182 | 302 | 598 | 602 | 243 | 351 |  |
| Col 55 X 3162 55 | 13.71 | 19.65 | 21.2 | 18.98 | 14.17 | 15.34 | 22.06 | 23 | 21.12 | 18.8033 |
|  | 226 | 234 | 580 | 158 | 261 | 352 | 169 | 552 | 232 |  |
| **Comparison between different groups** | | | | | | | | | | |
| Cross | | | | | | P Value | | Significance | | |
| 3162 40 X Col 40-3162 45 X Col 45 | | | | | | 0.54900 | | NS | | |
| 3162 40 X Col 40-3162 50 X Col 50 | | | | | | 0.00439 | | ** | | |
| 3162 40 X Col 40-3162 55 X Col 55 | | | | | | 0.25239 | | NS | | |
| 3162 45 X Col 45-3162 50 X Col 50 | | | | | | 0.34817 | | NS | | |
| 3162 45 X Col 45-3162 55 X Col 55 | | | | | | 0.99924 | | NS | | |
| 3162 50 X Col 50-3162 55 X Col 55 | | | | | | 0.70599 | | NS | | |
| Col 40 X 3162 40- Col 45 X 3162 45 | | | | | | 0.99810 | | NS | | |
| Col 40 X 3162 40- Col 50 X 3162 50 | | | | | | 0.99999 | | NS | | |
| Col 40 X 3162 40- Col 55 X 3162 55 | | | | | | 0.99996 | | NS | | |
| Col 45 X 3162 45- Col 50 X 3162 50 | | | | | | 0.99991 | | NS | | |
| Col 45 X 3162 45- Col 55 X 3162 55 | | | | | | 0.94928 | | NS | | |
| Col 50 X 3162 50- Col 55 X 3162 55 | | | | | | 0.99602 | | NS | | |
| 3162 40 X Col 40- Col 40 X 3162 40 | | | | | | 0.50437 | | NS | | |
| 3162 45 X Col 45- Col 45 X 3162 45 | | | | | | < 0.001 | | *** | | |
| 3162 50 X Col 50- Col 50 X 3162 50 | | | | | | < 0.001 | | *** | | |
| 3162 55 X Col 55- Col 55 X 3162 55 | | | | | | < 0.001 | | *** | | |

**Table S8: CO rates in the progeny of a cross between parents of CO detector line and wild type Columbia plants of different ages (CTL5.17)**

| **Cross** | **CO Rate (R)/Seeds per replicate (S)** | | | | **Mean CO Rate** | |
| --- | --- | --- | --- | --- | --- | --- |
|  | R1/S1 | R2/S2 | R3/S3 | |  |  |
| CTL5.17 40 X Col 40 | 4.42/475 | 5.39/334 | 4.9/354 | | 4.9033 | |
| Col 40 X CTL5.17 40 | 6.03/745 | 5.24/768 | 4.7/399 | | 5.3233 | |
| CTL5.17 45 X Col 45 | 3.88/1409 | 6.25/687 | 5.07/958 | | 5.0667 | |
| Col 45 X CTL5.17 45 | 6.08/704 | 5.93/661 | 4.78/364 | | 5.5967 | |
| CTL5.17 50 X Col 50 | 16.51/644 | 16.67/304 | 17.79/405 | | 16.99 | |
| Col 50 X CTL5.17 50 | 17.9/517 | 17.85/392 | 15.66/421 | | 17.1367 | |
| CTL5.17 55 X Col 55 | 19.54/477 | 12.05/354 | 16.02/278 | | 15.87 | |
| Col 55 X CTL5.17 55 | 18.65/252 | 16.1/354 | 14.78/421 | | 16.51 | |
| **Comparison between different groups** | | | | | | |
| Cross | | | | P Value | | Significance |
| CTL5.17 40 X Col 40-CTL5.17 45 X Col 45 | | | | 1.000 | | NS |
| CTL5.17 40 X Col 40-CTL5.17 50 X Col 50 | | | | 1.000 | | NS |
| CTL5.17 40 X Col 40-CTL5.17 55 X Col 55 | | | | 0.999 | | NS |
| CTL5.17 45 X Col 45-CTL5.17 50 X Col 50 | | | | 1.000 | | NS |
| CTL5.17 45 X Col 45-CTL5.17 55 X Col 55 | | | | 1.000 | | NS |
| CTL5.17 50 X Col 50-CTL5.17 55 X Col 55 | | | | 1.000 | | NS |
| Col 40 X CTL5.17 40- Col 45 X CTL5.17 45 | | | | 1.000 | | NS |
| Col 40 X CTL5.17 40- Col 50 X CTL5.17 50 | | | | 0.987 | | NS |
| Col 40 X CTL5.17 40- Col 55 X CTL5.17 55 | | | | 1.000 | | NS |
| Col 45 X CTL5.17 45- Col 50 X CTL5.17 50 | | | | 0.974 | | NS |
| Col 45 X CTL5.17 45- Col 55 X CTL5.17 55 | | | | 1.000 | | NS |
| Col 50 X CTL5.17 50- Col 55 X CTL5.17 55 | | | | 1.000 | | NS |
| CTL5.17 40 X Col 40- Col 40 X CTL5.17 40 | | | | <.001 | | *** |
| CTL5.17 45 X Col 45- Col 45 X CTL5.17 45 | | | | <.001 | | *** |
| CTL5.17 50 X Col 50- Col 50 X CTL5.17 50 | | | | <.001 | | *** |
| CTL5.17 55 X Col 55- Col 55 X CTL5.17 55 | | | | <.001 | | *** |

**Table S9: Line-wise male and female mean ratio comparisons**

| **CO line** | **Mean** | | **SEM** | |
| --- | --- | --- | --- | --- |
| CTL1.2 | 1.1875 | | 0.124466 | |
| CTL1.18 | 2.35 | | 0.409959 | |
| CTL2.4 | 1.1875 | | 0.124466 | |
| COL3-4/20 | 2.6875 | | 0.299819 | |
| 3158 | 2.335 | | 0.13404 | |
| CTL4.7 | 2.6775 | | 0.209821 | |
| 3162 | 1.2275 | | 0.194829 | |
| CTL15.17 | 3.1925 | | 0.216391 | |
| **Comparison between different CO lines** | | | | |
| **Cross** | | **P Value** | | **Significance** |
| CLT2.4-3158 | | <0.001 | | *** |
| CLT2.4-3162 | | 0.9976 | | Non-Significant (NS) |
| CLT2.4-CLT4.7 | | <0.001 | | *** |
| CLT2.4-CLT5.17 | | <0.001 | | *** |
| CLT2.4-Col3-4/20 | | <0.001 | | *** |
| CLT2.4-CLT1.18 | | <0.001 | | *** |
| CLT2.4-CLT1.2 | | 0.9853 | | NS |
| 3158-3162 | | <0.001 | | *** |
| 3158-CLT4.7 | | 0.4759 | | NS |
| 3158-CLT5.17 | | <0.001 | | *** |
| 3158-Col3-4/20 | | 0.4365 | | NS |
| 3158-CLT1.18 | | 1.0000 | | NS |
| 3158-CLT1.2 | | <0.001 | | *** |
| 3162-CLT4.7 | | <0.001 | | *** |
| 3162-CLT5.17 | | <0.001 | | *** |
| 3162-Col3-4/20 | | <0.001 | | *** |
| 3162-CLT1.18 | | <0.001 | | *** |
| 3162-CLT1.2 | | 1.0000 | | NS |
| CLT4.7-CLT5.17 | | 0.0514 | | NS |
| CLT4.7-Col3-4/20 | | 1.0000 | | NS |
| CLT4.7-CLT1.18 | | 0.5366 | | NS |
| CLT4.7-CLT1.2 | | <0.001 | | *** |
| CLT5.17-COL3-4/20 | | 0.0616 | | NS |
| CLT5.17-CLT1.18 | | <0.001 | | *** |
| CLT5.17-CLT1.2 | | <0.001 | | *** |
| COL3-4/20-CTL1.18 | | 0.4962 | | NS |
| COL3-4/20-CTL1.2 | | <0.001 | | *** |
| CTL1.18-CTL1.2 | | <0.001 | | *** |

**Table S10: Age-wise male and female mean ratio comparisons**

| **Age** | **Mean** | | **SEM** | |
| --- | --- | --- | --- | --- |
| 40 | 2.10125 | | 0.835728 | |
| 45 | 2.21 | | 0.856371 | |
| 50 | 2.2 | | 0.762234 | |
| 55 | 1.98875 | | 0.723196 | |
| **Comparison between different age groups** | | | | |
| **Age groups** | | **P Value** | | **Significance** |
| 40-45 | | 0.993 | | NS |
| 40-50 | | 0.995 | | NS |
| 40-55 | | 0.992 | | NS |
| 45-50 | | 1.000 | | NS |
| 45-55 | | 0.945 | | NS |
| 50-55 | | 0.952 | | NS |

**Table S11: Seed count for both male and females of different ages for the lines used**

| Age | CTL1.2 | CTL1.18 | CTL2.4 | COL3-4/20 | 3158 | CTL4.7 | 3162 | CTL5.17 |
| --- | --- | --- | --- | --- | --- | --- | --- | --- |
| 40 **♂** | 1548 | 1844 | 884 | 1803 | 2369 | 910 | 2049 | 1912 |
| 40 **♀** | 1765 | 2205 | 1490 | 1797 | 4076 | 1966 | 3362 | 1163 |
| 45 **♂** | 1586 | 2019 | 1791 | 3223 | 3224 | 2282 | 3248 | 1729 |
| 45 **♀** | 2417 | 2630 | 1363 | 4536 | 5028 | 2409 | 5076 | 3054 |
| 50 **♂** | 1115 | 1243 | 1181 | 2783 | 2707 | 1306 | 2335 | 1330 |
| 50 **♀** | 865 | 991 | 1738 | 2676 | 3129 | 1442 | 2347 | 1353 |
| 55 **♂** | 1425 | 1072 | 486 | 2290 | 3565 | 977 | 2764 | 1027 |
| 55 **♀** | 2550 | 1880 | 918 | 3533 | 5057 | 2694 | 4203 | 1109 |
